# Membrane Association and Functional Mechanism of Synaptotagmin-1 in Triggering Vesicle Fusion

**DOI:** 10.1101/2020.03.13.991588

**Authors:** Ramesh Prasad, Huan-Xiang Zhou

## Abstract

Upon Ca^2+^ influx, synaptic vesicles fuse with the presynaptic plasma membrane (PM) to release neurotransmitters. Membrane fusion is triggered by synaptotagmin-1, a transmembrane protein in the vesicle membrane (VM), but the mechanism is under debate. Synaptotagmin-1 contains a single transmembrane helix (TM) and two tandem C2-domains (C2A and C2B). The present study aimed to use molecular dynamics simulations to elucidate how Ca^2+^-bound synaptotagmin-1, by simultaneously associating with VM and PM, brings them together for fusion. While C2A stably associates with VM via two Ca^2+^-binding loops, C2B has a propensity to partially dissociate. Importantly, an acidic motif in the TM-C2A linker competes with VM for interacting with C2B, thereby flipping its orientation to face PM. Subsequently C2B can readily associate with PM via a polybasic cluster and a Ca^2+^-binding loop. These results delineate the functional process of fusion triggered by synaptotagmin-1.

## Introduction

At the synapse, fast synchronous release of neurotransmitters upon Ca^2+^ influx is essential for the integrity of neurotransmission. Neurotransmitter release results from fusion of synaptic vesicles with the presynaptic plasma membrane (PM). The fusion process is driven by the assembly of the trans-SNARE complex, comprising synaptobrevin tethered to the vesicle membrane (VM) and syntaxin and SNAP25 tethered to PM^1^. However, it is synaptotagmin-1 (Syt1), a transmembrane protein in the VM, that acts as the Ca^2+^ sensor and triggers the fusion of VM and PM^2^. How Syt1 carries out this function is still hotly debated^1, 3-6^. The present study aimed to use molecular dynamics (MD) simulations of full-length Syt1 associating with VM and PM to provide crucial missing information and develop a complete mechanism for the triggering of membrane fusion by Syt1.

Syt1 contains an N-terminal single transmembrane helix (TM; for tethering to VM), a disordered linker, and two cytoplasmic, Ca^2+^-binding C2 domains termed C2A and C2B (Fig. 1a and Supplementary Fig. 1). Each C2 domain folds into an eight-stranded β sandwich, and two loops (referred to as loop 1 and loop 3) at the top form the Ca^2+^-binding site (Fig. 1b-d)^7-9^. C2A binds three Ca^2+^ ions whereas C2B binds two. Upon Ca^2+^ binding, these loops penetrate into acidic membranes^10-13^. While phosphatidylserine (PS) is the major acidic lipid present in both VM and PM at >10% levels, the highly acidic phosphatidylinositol(4,5)-bisphosphate (PIP2; Supplementary Fig. 2) is also present in the inner leaflet of PM and reaches as much as 6% at the site of fusion^14, 15^. In addition to the Ca^2+^-binding loops, other sites of Syt1 have been recognized as important for binding with membranes or SNARE, including a conserved polybasic cluster (K_324_KKK_327_) on the side of C2B, and a conserved RR motif (residues 398-399) at the bottom of C2B. Moreover, the disordered linker harbors a conserved TM-proximal basic motif and a conserved C2A-proximal acidic motif (Fig. 1a).

**Figure 1.**
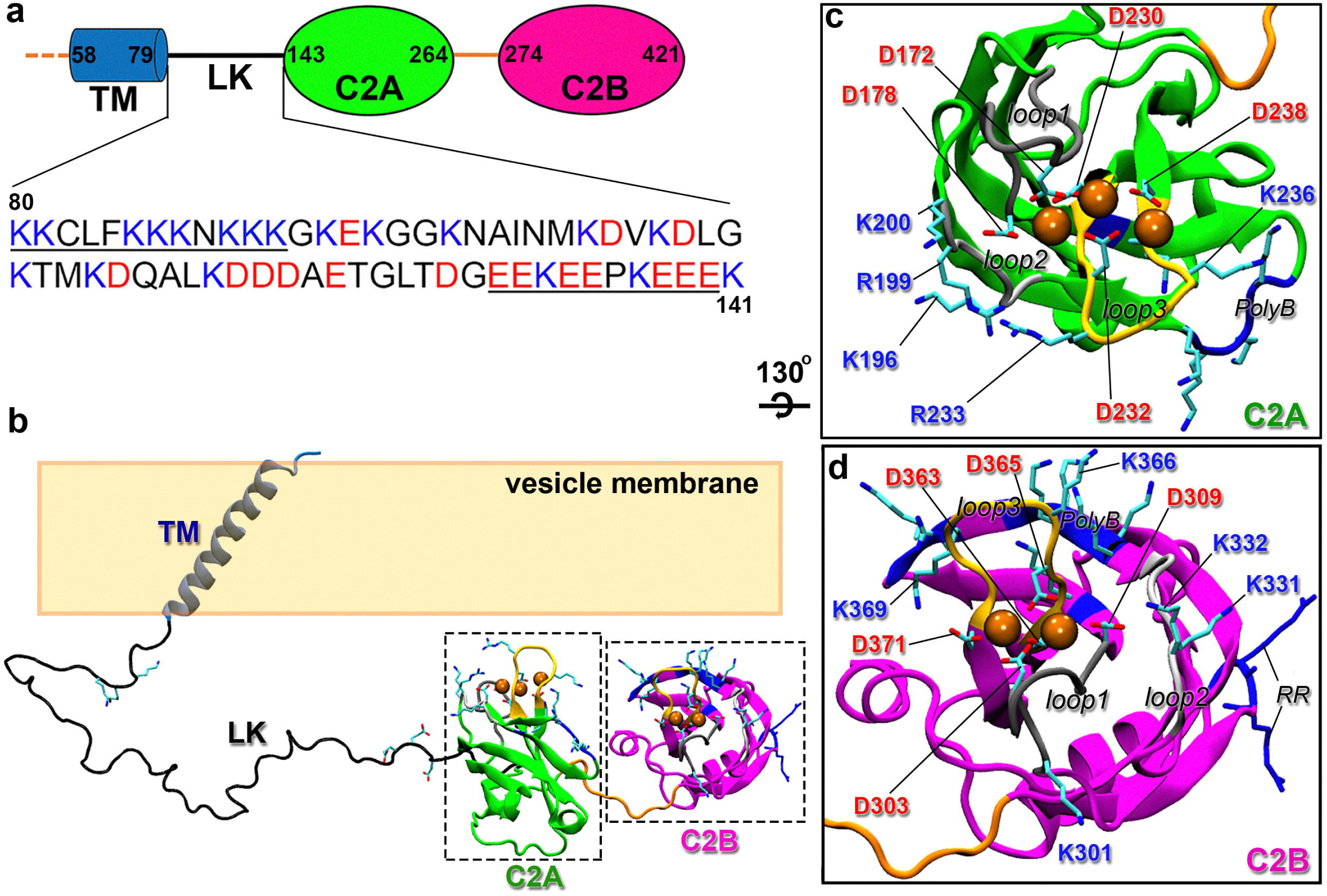
Sequence and domain structures of Syt1. (a) Syt1 domain decomposition, containing a transmembrane helix (TM), a disordered linker (LK), and two C2 domains (C2A and C2B). LK contains a basic motif and an acidic motif (underlined sequence). (b) Positioning of the domains of Ca^2+^-bound Syt1 relative to the vesicle membrane. Selected acidic and basic side chains in LK, C2A, and C2B are shown as sticks; bound Ca^2+^ ions are shown as spheres. (c) Enlarged view of C2A, highlighting the two Ca^2+^-binding loops (loops 1 and 3, with backbone in dark gray and yellow, respectively). Aspartates coordinating Ca^2+^ ions and basic residues in loop 1, loop3, loop 2 (backbone in light gray), and the polybasic cluster (polyB; backbone in blue) are shown as sticks. (d) Corresponding drawing for C2B. In addition to those with counterparts in C2A, the RR motif is shown with both the backbone and side chains in blue.

The identities of the interaction partners of the foregoing sites, the relevance of these interactions to membrane fusion, and the time sequence of these interactions during fusion are a mix of consensus and controversy. Using isolated C2A and C2B domains and membranes containing either PS or PIP2 as the only acidic lipid, Bai et al.^11^ obtained fluorescence intensity data to show that the Ca^2+^-binding loops of C2A penetrated PS-containing membranes only whereas those of C2B penetrated PIP2-containing membranes only, implying that C2A and C2B might be specialized in binding to VM and PM, respectively. The preference of C2A for PS-containing membranes has been confirmed by electron paramagnetic resonance (EPR)^16^, but supporting evidence for the preference of C2B for PIP2-containing membranes is less direct, especially in the context of a full-length Syt1 tethered to a PS-containing membrane^17^. There is evidence that the polybasic cluster of C2B contributes to its binding with PIP2-containing membranes^13, 16, 18-22^. It has also been suggested that the RR motif of C2B, at the end opposite to the Ca^2+^-binding loops, may also bind acidic membranes^4, 13, 23-25^. Both the polybasic cluster and RR motif have also been proposed to interact with SNARE^9, 22, 25, 26^, therefore raising the possibility that membranes and SNARE may compete for the same sites on Syt1. Wang et al.^22^ have contended that the RR-PS binding is weak and is replaced by RR-SNARE binding during fusion. It should be noted that C2A also has a conserved polybasic cluster (K_189_KKK_192_; Fig. 1c and Supplementary Fig. 1)^7^, but its role has been less studied^26-28^.

Based on its membrane binding properties, different groups have proposed that Syt1 may trigger fusion by bridging VM and PM^4, 13, 17, 20, 21, 23-25, 28-31^. The bridging can be achieved with C2A bound to VM and C2B bound to PM, or with C2B simultaneously bound to VM and PM, or with C2A bound to VM while C2B bound to both VM and PM. The bridging may simply shorten the distance between the two membranes, or reduce potential electrostatic repulsion between the acidic membranes, or direct the final step of the assembly of the SNARE complex. Syt1 has also been proposed to bend membranes for fusion^32-35^. In addition, there is evidence supporting the involvement of Syt1 in the docking and priming of vesicles prior to Ca^2+^ influx^1, 3, 4^.

Many of the fusion assays have used isolated C2 domains or the tandem C2 construct (termed C2AB). These constructs may miss crucial elements and therefore mislead mechanistic understanding^36^. For example, by tethering to VM via TM, cis binding of the C2 domains in full-length Syt1 is kinetically favored over trans binding to PM, which involves diffusion of the membranes toward each other^37^. This distinction between cis and trans binding is lost when using the C2AB construct. Moreover, the linker between TM and C2A has been found to be important. Noting the conserved TM-proximal basic and C2A-proximal acidic motifs, Lai et al.^38^ showed that charge inversion of either motif suppressed fusion. They suggested that electrostatic attraction kept the two motifs at a close distance, but a recent EPR study found that that was not the case^17^. In any event, Lee and Littlelon^39^ found that deletion of the linker abolished synchronous neurotransmitter release, while duplicating the linker maintained normal function. In addition, both switching the order of the two C2 domains and tethering of Syt1 to PM also abolished synchronous release. On the computational side, electrostatic modeling and MD simulations have only been done on isolated C2 domains on membranes^21, 35, 40, 41^ or the C2AB construct in water^42^.

Here, for the first time, we carried out MD simulations of full-length Syt1 to probe their membrane association. When tethered to VM (containing only PS as acidic lipids) by TM, C2A stably associates with VM via the two Ca^2+^-binding loops, but C2B has a propensity to partially dissociate. The dissociation is helped in part by the acidic motif in the disordered linker transiently interacting with the Ca^2+^-binding loops of C2B, thereby flipping it to potentially face PM. In comparison, C2A and C2B both stably associate with PM (containing both PIP2 and PS). When the C2B flipped from VM is placed near PM, C2B quickly associates with PM, hence bridging the space between the two membranes with C2A bound to VM and C2B bound to PM. This bridged configuration can facilitate the full assembly of the trans-SNARE complex, thereby accelerating membrane fusion.

## Results

We carried out MD simulations of the full-length Syt1 (FL; actually residues 53-418, without the lumenal N-terminus and missing the C-terminal three residues) as well as several shorter constructs: the disordered linker (LK; residues 80-141), FL with the C2 domains truncated (FLΔC2AB; residues 53-141), FL with C2B truncated (FLΔC2B; residues 53-264), and the two tandem C2 domains (C2AB; residues 143-418). These constructs were tethered to or on VM (composition: PS:PC:PE:cholesterol = 40:30:20:10), PM (PIP2:PS:PC:PE:cholesterol = 4:15:51:20:10 for inner leaflet), or both. We ran four replicate simulations (referred to as sim1 to sim4), each 1 μs long, for constructs (FL, FLΔC2B, FLΔC2AB, and LK) interacting with VM only; two replicate simulations, each 0.5 μs long, for a construct (C2AB) interacting with PM only; and one simulation (0.4 μs) for C2AB interacting with both VM and PM.

### The linker has minimal VM association and self-interaction

We first probed the extent to which the disordered linker associates with VM when Syt1 is tethered to this membrane via TM. According to the average distances 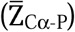 of C_α_ atoms from the plane of lipid phosphate atoms (Fig. 2a), only the very TM-proximal lysines are frequently in contact with VM. At the neutral residue N88, the linker starts to peel off from the membrane. This 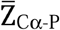 pattern for the first half of the linker holds regardless of the sequence truncations beyond the linker. Indeed, it remains true even for the isolated linker, except for the somewhat larger 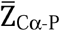 values for the first two lysines due to the lack of TM tethering. For most of the simulation time the other end of the linker does not come into contact with VM, due to electrostatic repulsion by the acidic motif (Supplementary Videos 1 and 2). However, 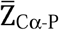 values do progressively reduce as the constructs get longer, due to the VM association of the C2 domains (see below). In line with the 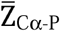 results, the average number of hydrogen bonds per residue with VM is high (> 0.5) for the first five lysines but decays rapidly after N88, with only occasional hydrogen bonds formed by lysines in the middle of the linker sequence (Fig. 2b). These results agree well with recent EPR data of Nyenhuis et al.^17^, showing that a spin label attached to residue 86 is in contact with lipids but those beyond (at 90, 95, 123, 136) are in aqueous environments.

**Figure 2.**
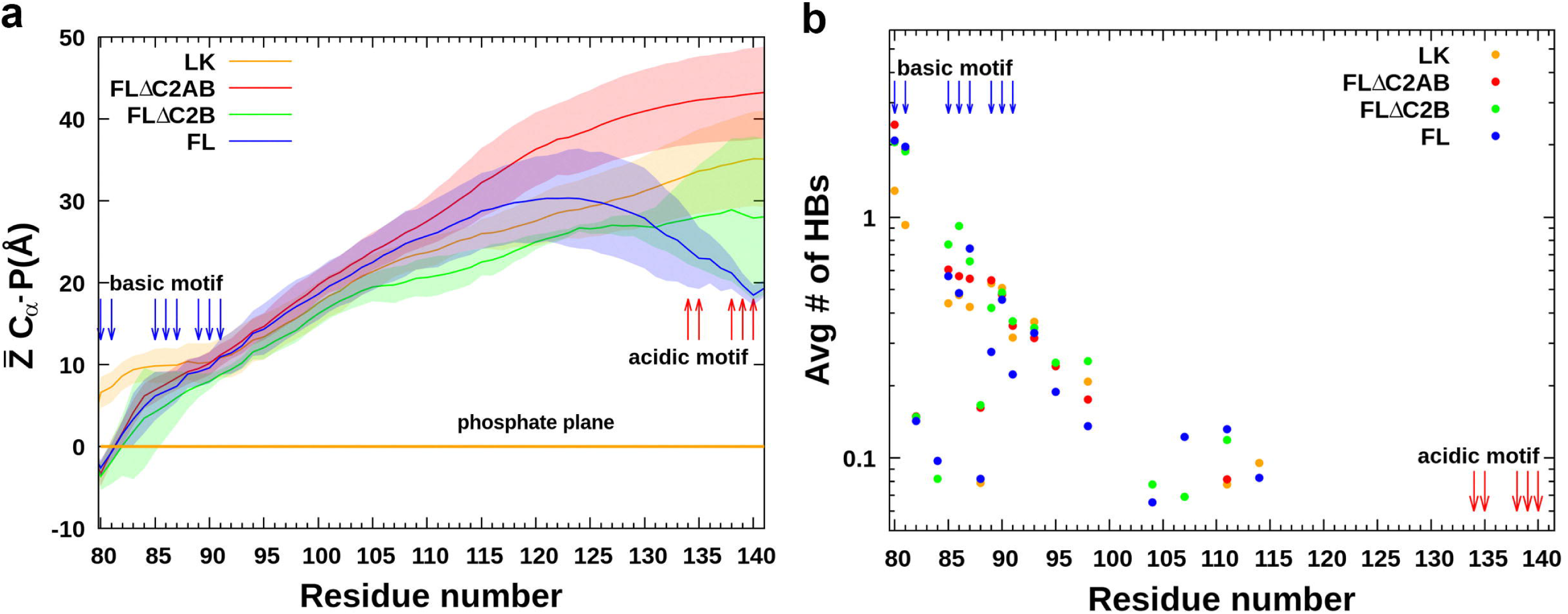
VM association of the disordered linker. (a) 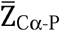, the average distance of each C_α_ atom from the phosphate plane. (b) The average number of hydrogen bonds (HBs) between a linker residue and lipids. Lysines and glutamates near the linker termini are indicated by blue and red arrows, respectively.

Lai et al.^38^ suggested that the basic and acidic motifs attract each other to compact the linker, but EPR data of Nyenhuis et al. showed that these motifs at most form transient interactions. Our MD simulations are consistent with the EPR data. For LK in solution, the basic and acidic motifs do frequently come into contact (Supplementary Video 3); the distances between the centers of geometry of K_85_KKNKKK_91_ and E_134_EPKEEE_140_ average at 27 A□ (Supplementary Fig. 3). Near VM, the contact between the basic and acidic motifs becomes much less frequent for LK, with the average distance between them increasing to 46 A□, because the basic motif is now engaged with VM and the acidic motif is repelled by acidic lipids (Supplementary Video 1). When tethered to VM via TM, any contact between the basic and acidic motifs is very transient, and their average distance further increases to around 65 A□ (Supplementary Video 2).

### C2A, but not C2B, stably associates with VM

In our FL-VM simulations, C2A stably associates with the membrane (Fig. 3a), with the number of residues in contact with lipids averaging around 12 (Supplementary Fig. 4a). In contrast, C2B on average has only about half as many residues forming contacts with lipids and sometimes dissociates from the membrane altogether (Supplementary Fig. 5a). We defined two parameters to characterize the position and orientation of each C2 domain relative to VM: the smallest displacement (Z) of any Cα atom in the domain from the phosphate plane and the angle (Θ) between the membrane normal and the vector from the Ca^2+^ ions to the center of the domain (Fig. 3b). The angle-displacement scatter plot of C2A shows two overlapping ensembles, one around Θ = 20° and Z = 2 A□ and the other around Θ = 50° and Z = −0.5 A□ (Fig. 3c), which will be called “straight” and “tilted”, respectively. In the simulations, C2A readily transitions between the straight and tilted configurations (Supplementary Fig. 4b). In the straight configuration (approximately 30% of all snapshots), VM association of C2A is stabilized by contacts with lipids by the three loops at the top; in the tilted configuration (approximately 70% of all snapshots), the polybasic cluster, in which we now include K182, participates in interacting with lipids, and meanwhile loop 1 becomes somewhat less important (Fig. 3d). Approximately the same number of C2A residues form contacts with VM in either the straight or tilted configuration (Supplementary Fig. 4). Overall, the three loops and the polybasic cluster of C2A provide nearly all the stabilization of VM association (Fig. 4a).

**Figure 3.**
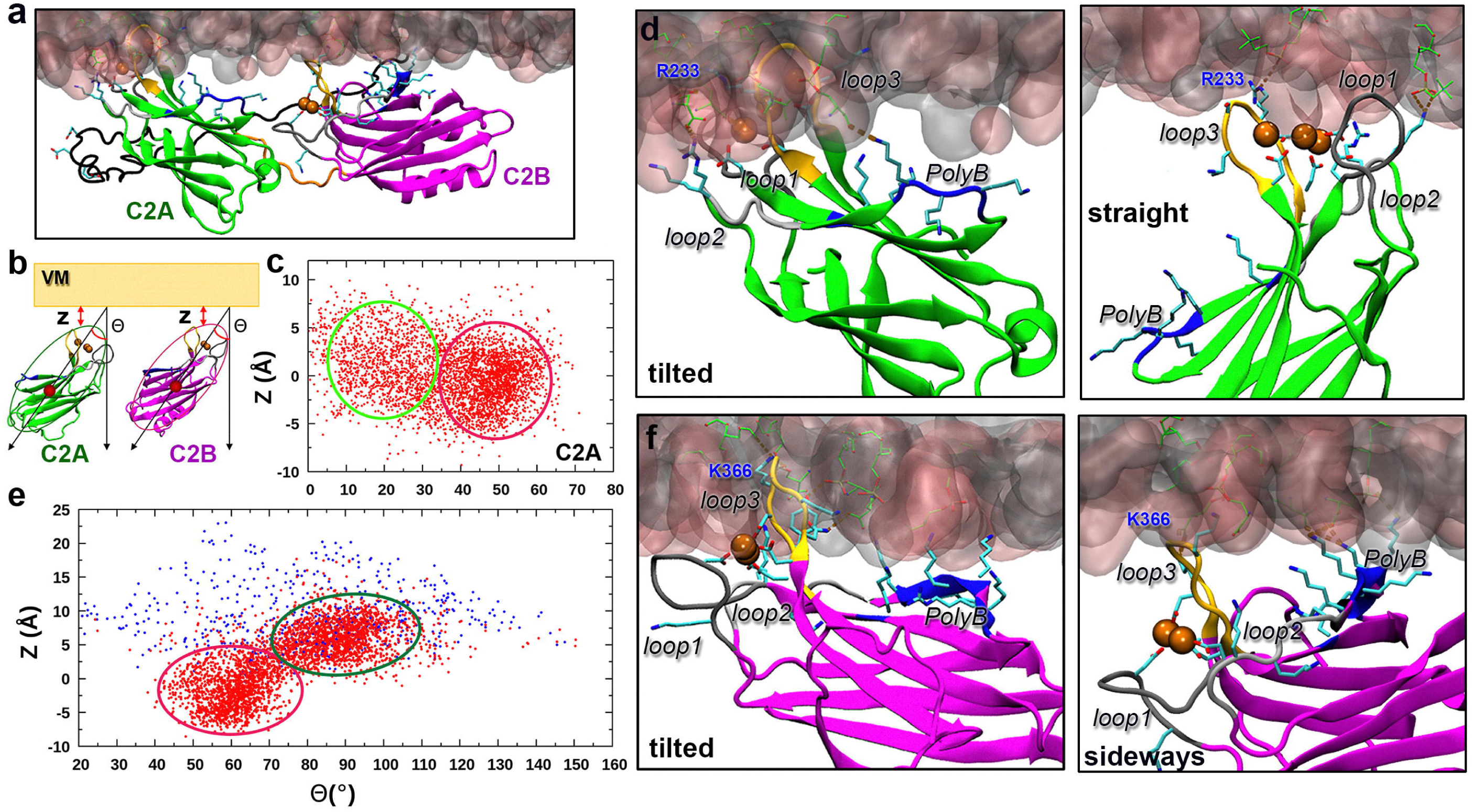
VM-associated configurations of C2A and C2B in FL simulations. (a) A snapshot from the FL-VM simulations. (b) Illustration of the smallest displacement (Z) of any Cα atom in a C2 domain from the phosphate plane and the tilt angle (Θ) of the domain. (c) Scatter plot of C2A Θ and Z collected from the simulations. Red and light green ovals indicate ensembles in the tilted and straight configurations, respectively. (d) Left: enlarged view of C2A in the tilted configuration shown in (a). Right: C2A in the straight configuration, from a different snapshot. R233 in loop 3 is labeled. (e) Scatter plot of C2B Θ and Z. Red and blue dots are from snapshots where C2B contacts and does not contact VM, respectively; red and dark green ovals indicate ensembles in the tilted and sideways configurations, respectively. (f) Right: enlarged view of C2B in the sideways configuration shown in (a). Left: C2B in the tilted configuration, from a different snapshot. K366 in loop 3 is labeled.

**Figure 4.**
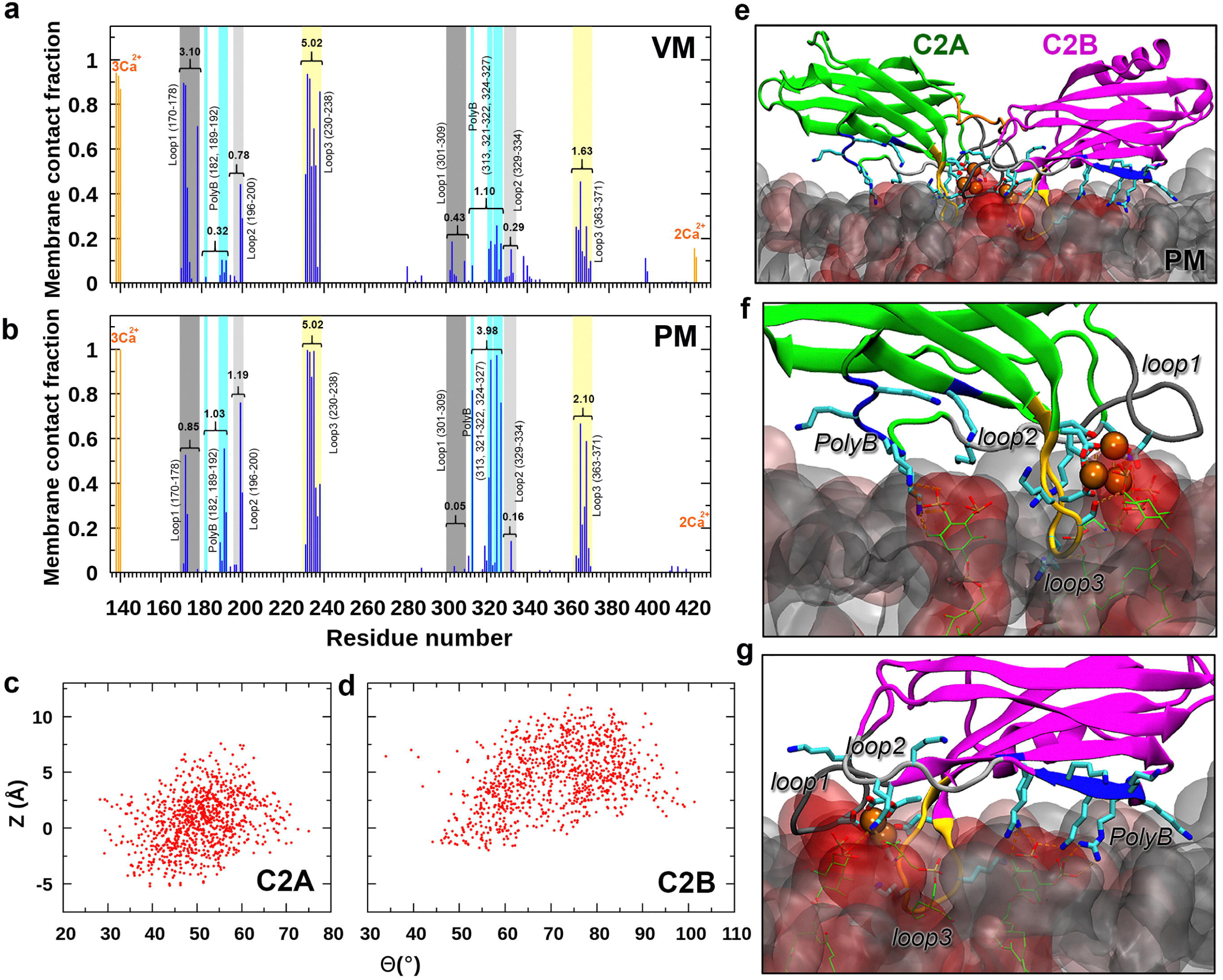
Comparison of VM- and PM-association of C2AB. (a) Fractions of time when individual C2AB residues form membrane contacts in the FL-VM simulations. The total membrane-contact numbers of different motifs (e.g., loop 3) are indicated. (b) Counterparts for the C2AB-PM simulations. (c) Scatter plot of C2A Θ and Z collected from the C2AB-PM simulations. (d) Scatter plot of C2B Θ and Z. (e) A snapshot from the C2AB-PM simulations. (f) Enlarged view of C2A, showing PM interactions. (g) Enlarged view of C2B.

In the dissociated snapshots (13.5% of total), Θ and Z of C2B are randomly scattered over wide ranges (21°-148° in Θ and 2.0-24.0 A□ in Z; Fig. 3e); below we will revisit C2B partial dissociation from VM. On the other hand, the VM-associated snapshots form two dense ensembles, one centered around Θ = 60° and Z = −2 A□ and the other around Θ = 90° and Z = 6 A□. We refer to these two types of configurations as tilted and sideways. In the tilted configuration (approximately half of the snapshots with VM contacts), loop 3 of C2B penetrates inside the phosphate plane, and there are opportunities for VM engagement by loops 1 and 2 and especially by the polybasic cluster, which we now widen to include K313, K321, and R322 (Fig. 3f, left panel). In the sideways configuration (approximately half of the snapshots with VM contacts), mostly it is the polybasic cluster, followed by loop 3, that interacts with lipids (Fig. 3f, right panel), but the membrane association is not very stable and C2B tends to roll on its side. Consequently other residues, including R281, K288, Y338, N340, and the RR motif (see below), also participate in lipid interactions occasionally. The number of C2B residues forming contacts with VM in the sideway configuration is less than that in the tilted configuration (averaging 3 and 8, respectively; Supplementary Fig. 5a). Overall, loop 3 and the polybasic cluster make the dominant contributions to C2B-VM association (Fig. 4a).

The more stable VM association of C2A relative to C2B seen in our MD simulations (Fig. 4a) is supported by several experimental studies. Using isolated C2 domains (Ca^2+^-bound), Hui et al.^43^ found that C2A complexes with PS membranes were much stronger than C2B complexes. The same study also found that a KAKA mutation, i.e., mutation of K326 and K327 in C2B to alanine, significantly weakened C2B-PS binding. Li et al.^19^ likewise obtained a destabilizing effect for the KAKA mutation on C2AB-PS binding, and an even greater effect for mutating R233 in C2A loop3 to glutamine, but a minimal effect for a corresponding mutation of the C2B loop 3 residue K366. The destabilizing effect of the KAKA mutation is in line with the significant involvement of the polybasic cluster of C2B in VM association in our simulations (Figs. 3f and 4a). The greater destabilization of the R233Q mutation validates our simulation result that loop 3 of C2A contributes the largest number of VM-contacting residues among all the motifs that potentially interact with VM (Fig. 4a). Lastly, the contrasting effects of the R233Q and K366Q mutations provide direct confirmation of the different extents of VM engagement by the loop 3’s of the two C2 domains in our simulations. Like its counterpart in C2A, loop 3 in C2B also contributes the largest number of VM-contacting residues within C2B, but the number of those residues is three times lower (5.02 for C2A vs. 1.63 for C2B). Of the two residues mutated by Li et al., R233 is the second most frequent VM-contacting residue in C2A (barely exceeded by the neighboring, Ca^2+^-coordinating D232) (Figs. 4a and 3d), while K366 is the runaway number one VM-contacting residue in C2B (Fig. 4a and 3f). Still, R233 is two times more likely to contact VM than K366.

In contrast to the stable VM association of C2A in FL, upon deletion of C2B, C2A has a tendency to dissociate from VM (Supplementary Fig. 6a). Dissociation occurs in 36% of all the snapshots. In the non-dissociated snapshots, approximately 80% are in the tilted configuration and the other 20% are in the sideways configuration (Supplementary Fig. 6b, c). Therefore the presence of a C2B that is not even stably associated with VM can still strengthen the association of C2A with VM. This stabilization also has direct experimental support. For example, Hui et al.^43^ found that the C2AB construct bound to PS membranes much more strongly than the isolated C2A domain. C2B provided stabilization even when its PS binding ability was reduced by loop 3 mutations that interfered with Ca^2+^ binding, although the extent of stabilization was commensurately reduced. Likewise EPR data of Herrick et al.^12^ suggested that C2AB positioned more deeply into PS membranes than the isolated C2A domain. Lastly kinetic measurements of Tran et al.^44^ showed that C2AB dissociated from PS membranes much more slowly than the isolated C2A domain. The stabilization of C2B can be rationalized theoretically by its effect in increasing the local concentration of C2A^45^.

### C2B stably associates with PM due to polyB-PIP2 interactions

Our C2AB-PM simulations show that both C2A and C2B stably associate with PM, with both adopting a tilted configuration (Fig. 4b-g). For C2A, the PM-associated tilted configuration is very similar to the VM-associated one (compare Fig. 3c, region outlined by a red oval, and Fig. 4c; also compare Fig. 3d, left panel, and Fig. 4f). However, C2B becomes more tilted, from centering around 60° in the VM-associated tilted configuration to around 70° when associated with PM (compare Fig. 3e, region outlined by a red oval, and Fig. 4d; also compare Fig. 3f, left panel, and Fig. 4g). This change in the C2B tilt angle comes from much greater participation of the polybasic cluster in interacting with PM (Fig. 4b compared with Fig. 4a), and is directly supported by EPR data of Kuo et al.^13^ The PM interactions of C2B is overwhelmingly with PIP2 (Fig. 4g).

Although C2A stably associates with PM in the C2AB construct, it should be recognized that, in the context of FL Syt1 that is tethered to VM via TM, C2A association with VM is favored over PM both kinetically^37^ and thermodynamically^45^, because of the TM-C2A linkage. Indeed, recent EPR data of Nyenhuis et al.^17^ demonstrated that C2A in FL preferentially associated in cis with PS-containing membranes even when PIP2-contaning membranes were present in trans.

### Interactions with linker acidic motif stabilize C2B when partially dissociated from VM

We more closely investigated why C2B in FL would partially dissociate from VM. The snapshot at 218 ns in sim1 where C2B has the largest tilt angle, of 150.4° (Supplementary Fig. 5b), meaning that this domain is flipped to have its bottom facing VM, caught our attention. In this snapshot, the Ca^2+^-binding loops at the top face the acidic motif of the TM-C2A linker, while the RR motif at the bottom interacts with VM (Supplementary Fig. 5c). We therefore looked for interactions between the acidic motif and the Ca^2+^-binding loops of C2B, and found that they started to contact at 152 ns. Fig. 5a presents their interactions at 193 ns of sim1. A video showing the simulation from 150 to 225 ns (Supplementary Video 4) gives the appearance that the contact with the acidic motif tips over C2B.

**Figure 5.**
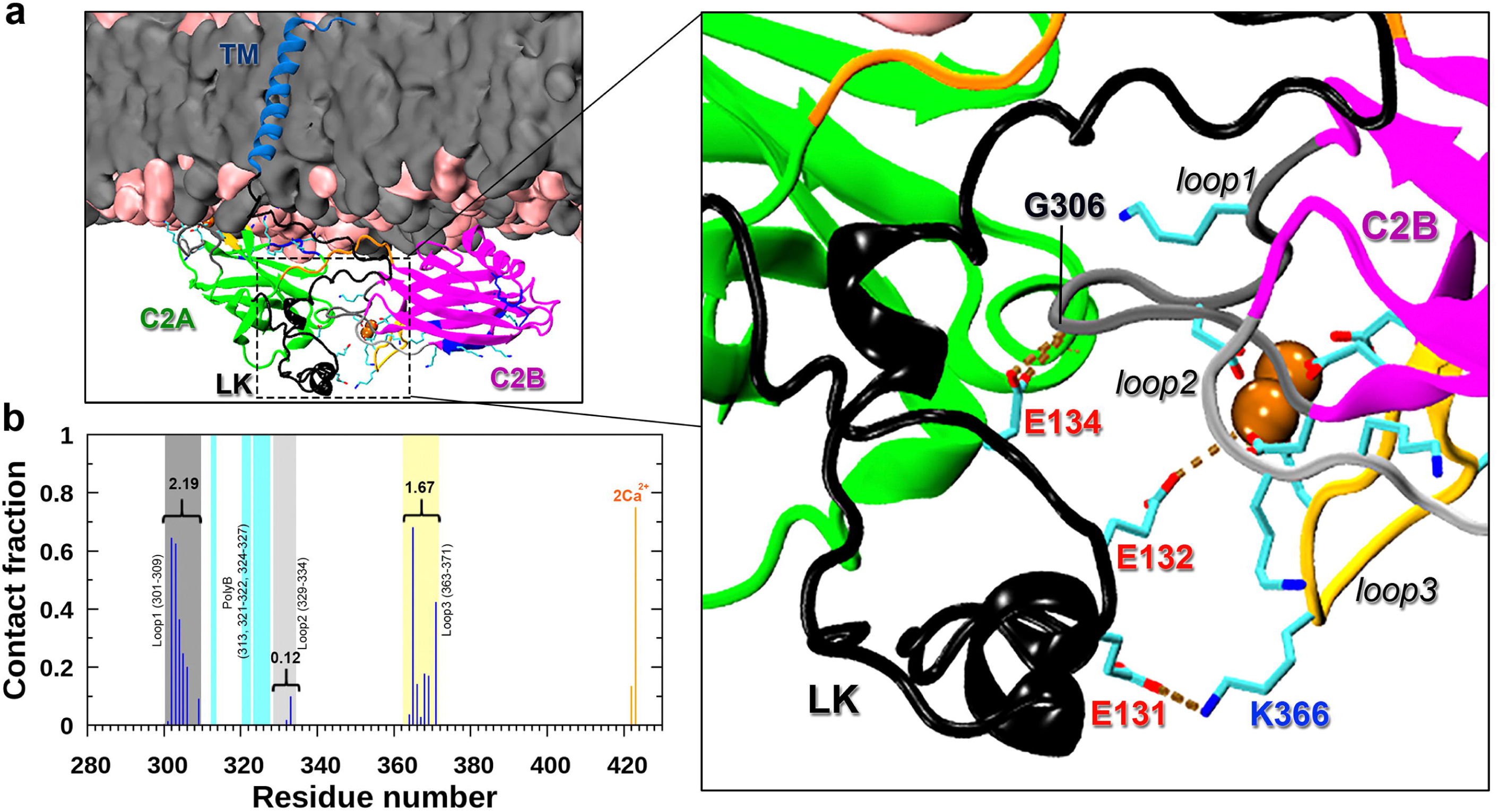
Interactions of the linker acidic motif with C2B Ca^2+^-binding loops. (a) Snapshot at 193 ns of FL-VM sim1. Enlarged view illustrates the interactions of Glu131, Glu132, and Glu134 interacting with loops 1 and 3 and a Ca^2+^ ion in C2B. (b) The fractions of snapshots where individual C2B residues form contacts with acidic residues in the linker, among all snapshots where at least one such contact is formed.

Contact of C2B with the acidic motif is actually quite prevalent, occurring in 10% of all snapshots in the FL-VM simulations. Nearly all the interactions of the acidic motif are with loop 1 and loop 2 of C2B (Fig. 5b). There is little chance for the acidic motif to interact with the polybasic cluster, in line with EPR data of Nyenhuis et al.^17^ It thus seems that the acidic motif of the TM-C2A linker competes with VM for C2B interaction, stabilizing C2B when it is partially dissociated from VM. The RR motif of C2B may also play a minor role in this regard.

### C2B released from VM readily associates with PM

To fully mimic the physiological situation of membrane fusion, we placed PM next to the C2B that was partially dissociated from VM (Fig. 6a; see Supplementary Video 5 for the entire simulation). While C2A remains stably associated with VM, C2B quickly (within ∼50 ns) approaches PM and changes its tilt angle to round 76° (Fig. 6b), close to the value typical of the stably bound configuration of C2B in the C2AB-PM simulations (Fig. 4d). In the next 230 ns or so, lipids rearrange around C2B and eventually PIP2 becomes the dominant lipid molecules interacting with the polybasic cluster, which drive up the total number of membrane contacts (Fig. 6c). In the remaining 120 ns, C2B is stably associated with PM (Fig. 6d).

**Figure 6.**
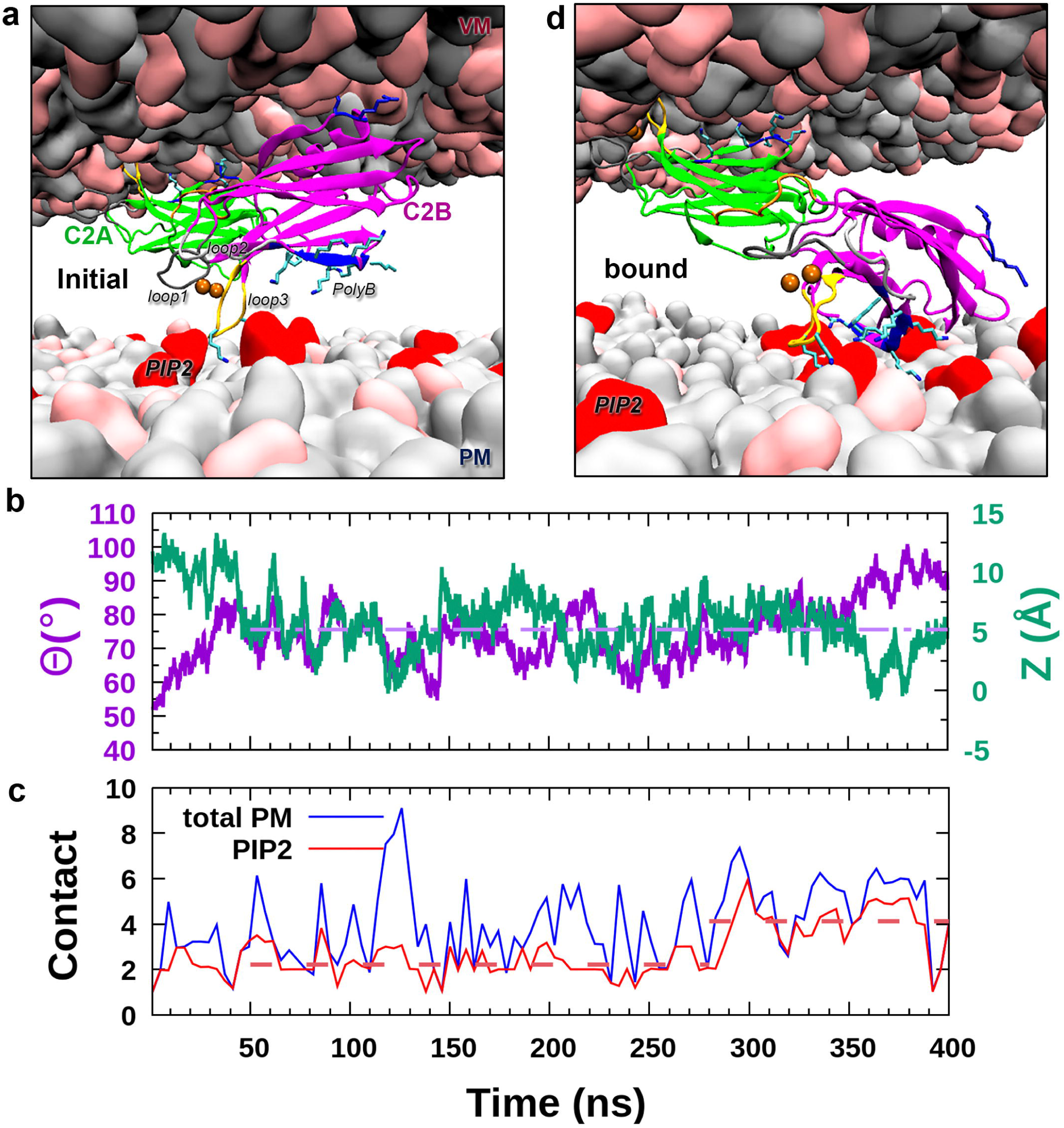
PM association of C2B released from VM. (a) Initial snapshot where C2B released from VM is placed near PM. (b) Time traces of Θ and Z in the simulation of C2AB associating with two membranes. A horizontal dashed line indicates the average Θ during 50 to 400 ns of the simulation. (c) Time trace of C2B contacts with PIP2 molecules or with all PM lipids. Horizontal dashed lines indicate the average C2B-PIP2 contact number from 50 to 280 ns and that for the remainder of the simulation. (d) A snapshot where C2B becomes stably associated with PM, showing the polybasic cluster and loop 3 (side chains shown as sticks) interacting with PIP2 molecules (shown as red surface).

## Discussion

Through extensive molecular dynamics simulations of FL Syt1 and several shorter constructs on both VM and PM, we have gained unprecedented information and insight on the membrane association and conformational transition that are likely to be relevant for triggering of membrane fusion by Syt1. When associated with either VM or PM, Syt1 is highly dynamic, with multiple configurations (e.g., tilted and straight for C2A) and each comprising a broad ensemble of poses, and sometimes C2B even dissociates from VM. Given this dynamic nature, molecular dynamics simulations are uniquely able to characterize the biophysical properties of the Syt1-membrane system. Still, our simulation results are directly supported by multiple experimental studies, including those reporting EPR data, membrane binding assays, and kinetic measurements.

The most interesting finding from the molecular dynamics simulations is that the conserved acidic motif in the TM-C2A linker can compete with VM for interacting with the Ca^2+^-binding loops of C2B, thereby flipping its orientation toward PM (Fig. 7, top row). A speculation that the acidic motif may help compact the linker by interacting with the upstream basic motif^38^ turned out to be inconsistent with subsequent EPR data.^17^ Nevertheless charge inversion of the acidic motif suppressed membrane fusion.^38^ Moreover, deletion of the linker abolished synchronous neurotransmitter release while duplicating the linker had no effect.^39^ These observations support the importance of the acidic motif, and interactions with the Ca^2+^-binding loops of C2B can explain this importance.

**Figure 7.**
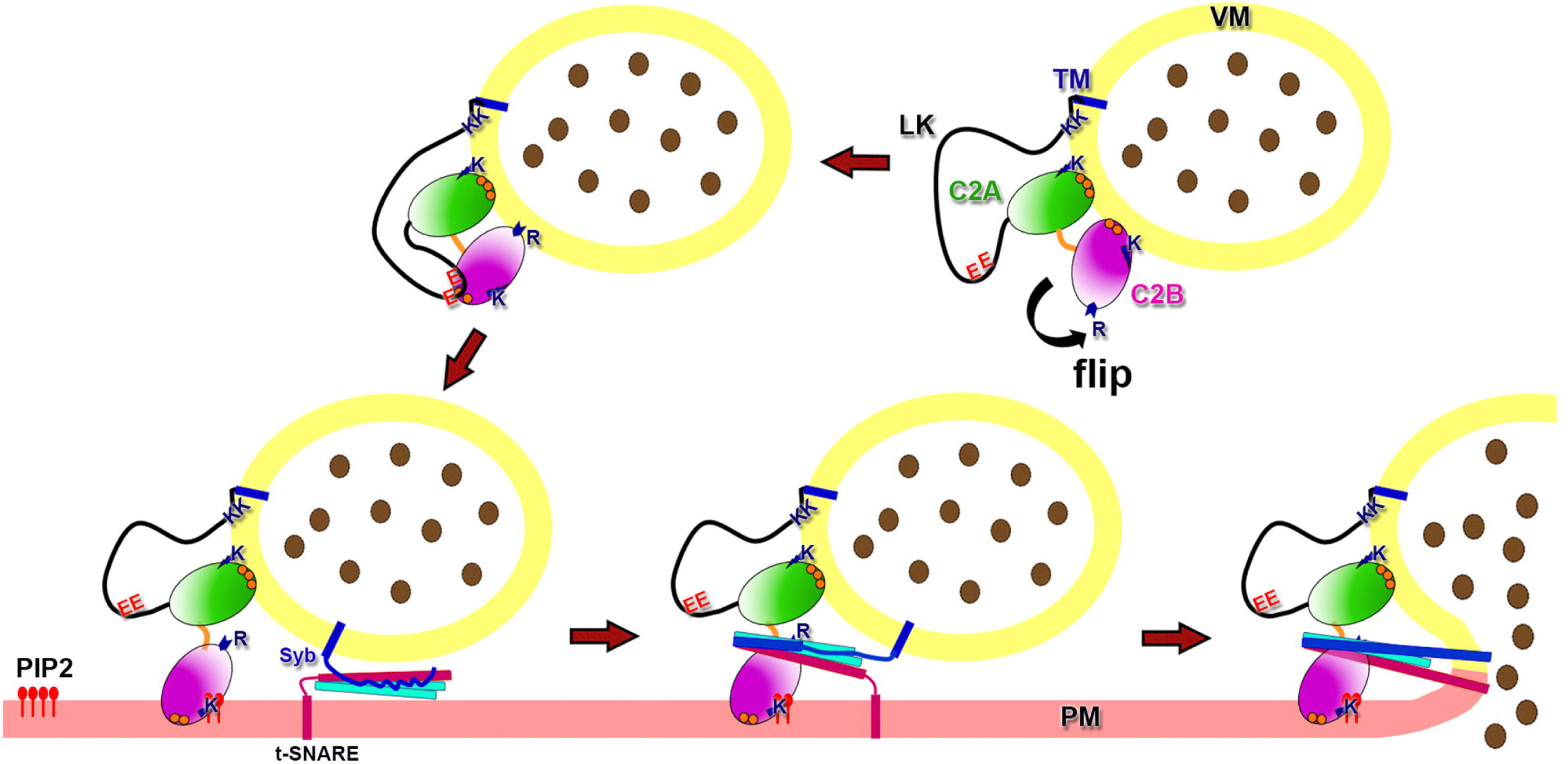
Proposed model for the triggering of membrane fusion by Syt1. By Interacting with the Ca^2+^-binding loops, the linker acidic motif flips C2B from VM-facing to PM-facing. C2B then quickly associates with PM, providing a platform for the complete assembly of the trans-SNARE complex.

Our simulations further showed that, when flipped from VM, C2B quickly associates with PM (Fig. 7, bottom row). Syt1 now bridges the space between the two membranes, with C2A bound to VM and C2B bound to PM. In this bridged configuration, C2B, especially through its RR motif,^9^ presents a platform for the trans-SNARE complex to complete its assembly. Membrane fusion then finally ensues. Our bridging model, in contrast to those relying on C2B simultaneously binding to both VM and PM, explains why it is important to preserve the order of C2A followed by C2B along the amino-acid sequence and to tether Syt1 to VM, as both switching the order of the two C2 domains and tethering of Syt1 to PM abolished synchronous release of neurotransmitters.^39^

While the present work has focused on Ca^2+^-bound Syt1, it will be interesting to apply molecular dynamics simulations to characterize the interactions of Syt1 prior to Ca^2+^ influx. Likewise it would be fruitful to compare and contrast Syt1, the primary Ca^2+^ sensor in synchronous neurotransmitter release, with other synaptotagmin isoforms, such as Syt7,^41, 44^ which plays a prominent role in asynchronous neurotransmitter release^46^. Lastly molecular dynamics simulations such as presented here can also help understand the lipid transport mechanisms of extended synaptotagmins, which are tethered to the endoplasmic reticulum and associate with PM via C2 domains^47^.

## Computational Methods

### System preparations

Two types of membranes (VM and PM) were generated using the CHARMM-GUI membrane builder server^48^. Each leaflet was composed of 400 lipids (see Supplementary Fig. 2 for molecular structures of the different types of lipids used). VM was symmetric, with the following composition: 40% DOPS, 30% DOPC, 20% DOPE, and 10% cholesterol. PM was asymmetric, with the following composition for the inner leaflet: 4% PIP2, 15% DOPS, 51% DOPC, 20% DOPE, and 10% cholesterol. For the outer leaflet, PIP2 was eliminated and made up by DOPC.

The Syt1 sequence was that from rat (NCBI protein sequence entry NP_001028852.2), which was the one studied in many experimental papers. Residues W58…C79 (corresponding to TM) was modeled as an α-helix, whereas upstream residues I53-P57 and downstream residues K80-K141 (corresponding to LK) were modeled as extended. The structure of C2AB was from protein data bank (PDB entry) 5KJ7 (chain E; residues K141-A418)^49^, but with three Ca^2+^ ions bound to C2A based on superposition to PDB entry 1BYN^8^. TM was inserted into VM manually in VMD^50^, and clashed lipids were removed.

### Molecular dynamics simulations

Amber ff14SB,^51^ Lipid17^52^, and TIP4PD^53^ (https://github.com/ajoshpratt/amber16-tip4pd) were used for modeling the proteins, lipids, and water, respectively. Lipid17 did not have parameters for PIP2, which we produced using the general Amber force field (GAFF)^54^. Specifically, GAFF supplied all parameters except for atomic charges, which we obtained by restrained fitting to the quantum mechanical (Gaussian 16 at the HF*/*6–31G* level) electrostatic potential. For LK in solution, the simulation box had dimensions of 100 x 100 x 70 A□^3^, allowing at least 12 A□ of solvent between the solute and the nearest box boundary. For LK and FLΔC2AB on VM, the box dimensions were 172 x 172 x 160 A□^3^; for FLΔC2B and FL, the dimensions were increased to 172 x 172 x 200 A□^3^. The box dimensions for C2AB on PM were similar, at 173 x 173 x 190 A□^3^. All the systems were neutralized by replacing water with K^+^ and Cl^-^, and the final KCl concentration was 150 mM. The solvation was done using the leap module of AMBER17 tools.

The systems were equilibrated in NAMD^55^. For those containing either VM or PM, a CHARMM-GUI recommended protocol^48^ was largely followed. After 10,000 cycles of energy minimization, each system was simulated at constant NVT with a timestep of 1 fs for 50 ps, with the protein and lipid headgroups restrained (the strength of lipid restraints was reduced to half in the second half of the simulation). Subsequently the simulation switched to constant NPT, initially (for 25 ps) with the lipid restraints further reduced by half but thereafter (for 600 ps) with the lipid restraints removed all together and the timestep increased to 2 fs. All the while the protein restraints were gradually reduced. The system was last equilibrated at constant NPT for 8 ns without any restraint.

Long-range electrostatic interactions were treated by the particle mesh Ewald method^56^, with non-bonded cutoff at 12 Å. All bonds connected to hydrogens were constrained using the SHAKE algorithm^57^. The temperature was maintained at 303 K using the Langevin thermostat with a damping coefficient of 1 ps^-1^, and the pressure was maintain at 1 atm using the Langevin piston method^58^.

The simulations for all the VM systems (and for LK in solution) were continued using *pmemd.cuda* on GPUs^59^. For each system, four replicate production runs were generated with different random seeds and each was extended to 1000 ns. For the PM system, two replicate productions runs were collected in NAMD, each for 500 ns. Snapshots were saved every ns.

For C2AB in the presence of both VM and PM, the system was prepared by taking C2AB and VM from a snapshot in the FL-VM simulations (at 222 ns of sim1) and PM from the C2AB-PM simulations. The inner phosphate planes of VM and PM were separated by 60 A□. The two membranes were trimmed down to approximately 280 lipids per leaflet, and the dimensions of the simulation box were 140 x 160 x 218 A□^3^. The system was energy minimized for 5,000 cycles, and equilibrated at constant NVT for 50 fs with a timestep of 1 fs, and with both the protein and lipids fixed. The production run then continued for 400 ns with a timestep of 2 fs. Snapshots were saved every 100 ps.

## Acknowledgments

This work was supported by National Institutes of Health Grant GM118091.

## Author contributions

H.X.Z. and R.P. designed the research. R.P. performed the research and analyzed the data.

H.X.Z. wrote the manuscript.

