## Supplementary Figures for "Membrane Association and Functional Mechanism of Synaptotagmin-1 in Triggering Vesicle Fusion"

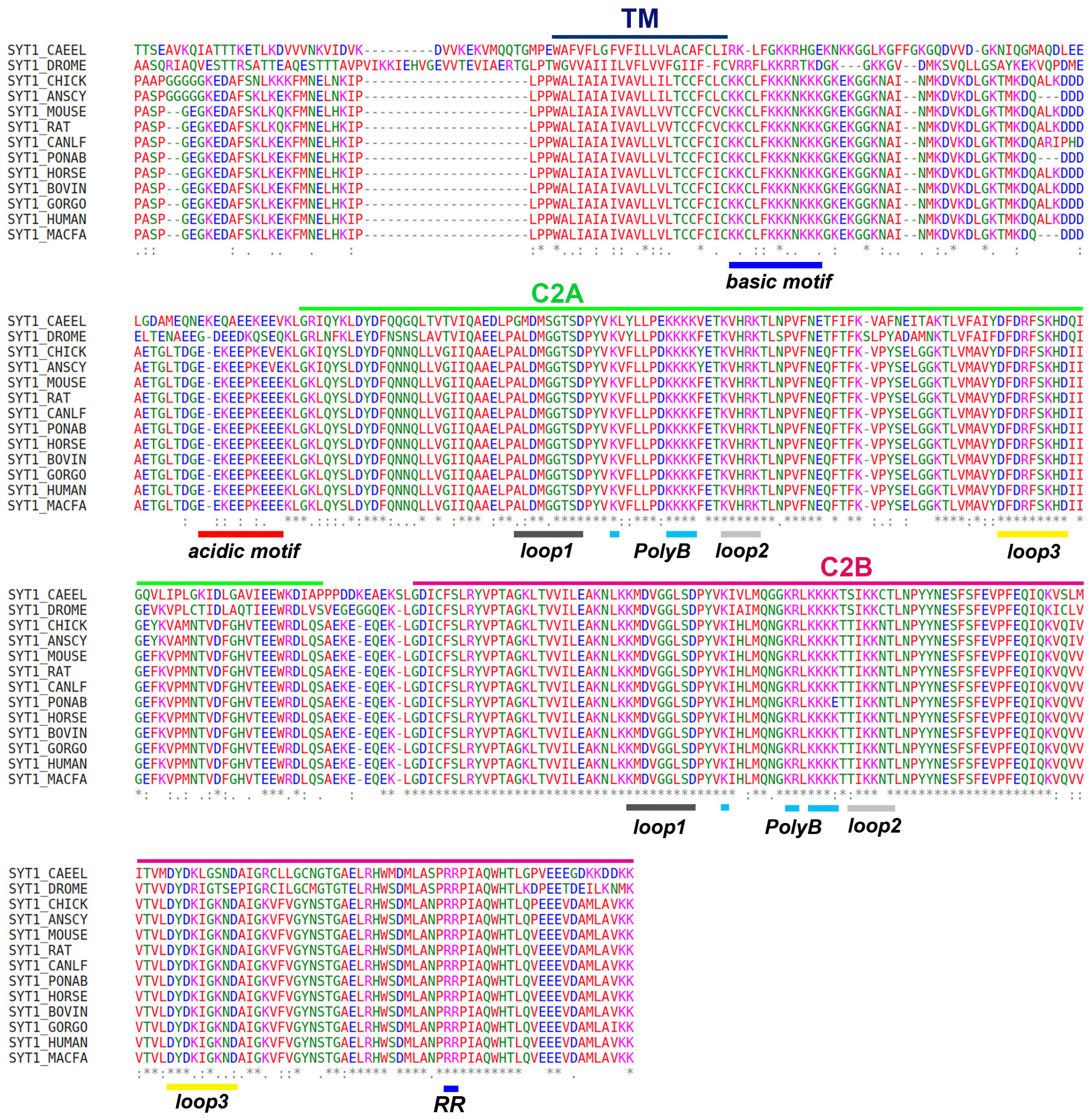


**Supplementary Figure 1. Sequence alignment of Syt1s from 13 organisms.** The basic and acidic motif in the TM-C2A linker, three loops (loops 1-3) and the polybasic cluster (PolyB) in each of the C2 domain, and the RR motif in C2B are indicated by thick line segments.

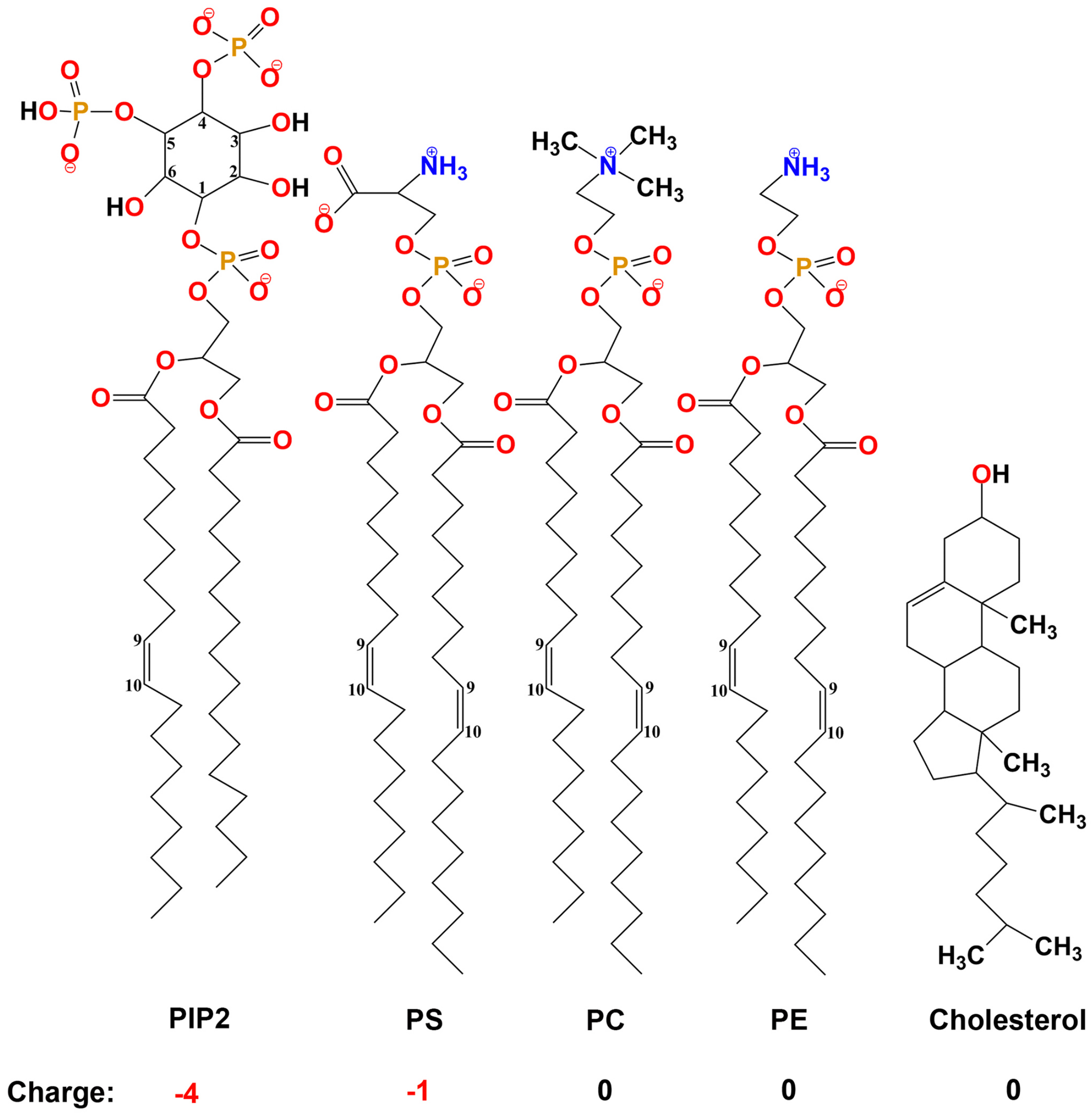


**Supplementary Figure 2. Molecular structures of the lipids in the present study.**

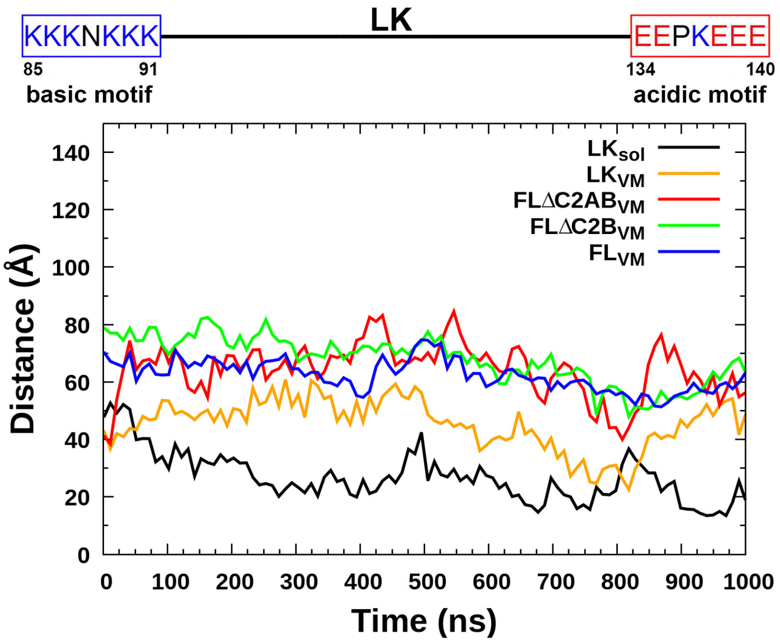


**Supplementary Figure 3. Time traces of the distance between a basic motif (residues 85-91) and an acidic motif (residues 134-140), in simulations of FL and three shorter constructs on VM as well as of LK in solution.** Each time trace represents the average of four replicate simulations.

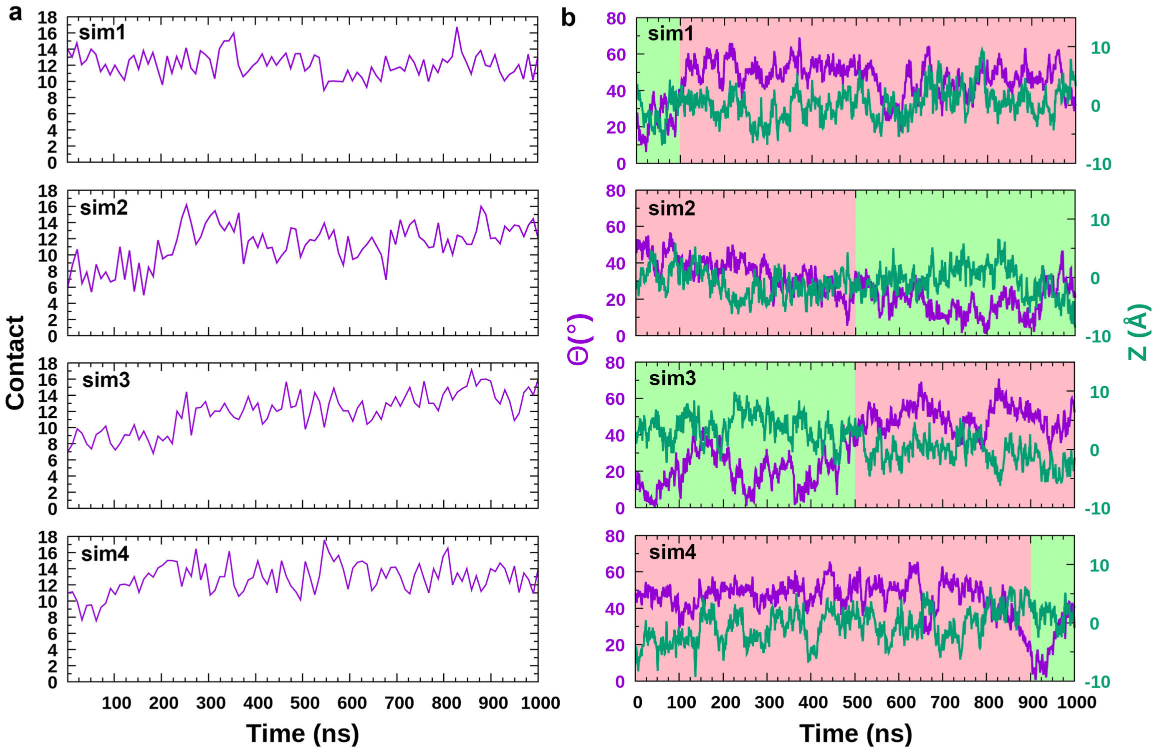


**Supplementary Figure 4. Membrane contacts and configurations of C2A in the FL-VM simulations.** (a) Time traces of the total VM-contact numbers. (b) Time traces of Θ and Z. Tilted and straight configurations are shaded in red and green, respectively.

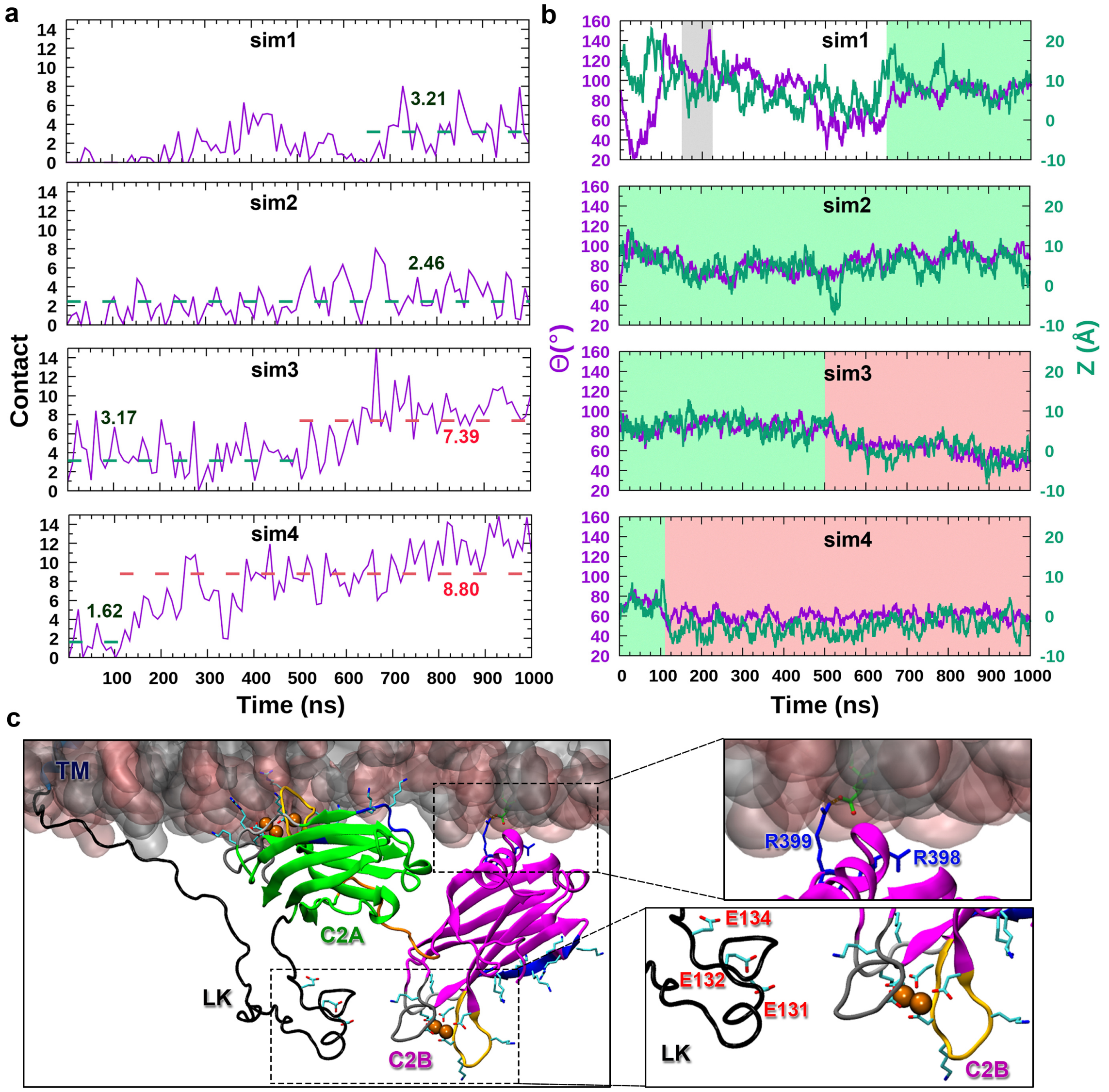


**Supplementary Figure 5. Membrane contacts and configurations of C2B in the FL-VM simulations.** (a) Time traces of the total VM-contact numbers. Horizontal dashes indicate average contact numbers in a tilted or sideways configuration during a simulation. (b) Time traces of Θ and Z. Tilted and sideways configurations are shaded in red and green, respectively. The segment shaded in gray in sim1 is where C2B is tipped over by interactions with the linker acidic motif; see Supplementary Video 4. (c) Snapshot at 218 ns in sim1, where C2B is flipped so its bottom faces VM. Enlarged views show that the $Ca^{2+}$-binding loops of C2B face the linker acidic motif, and the RR motif at the bottom interacts with VM.

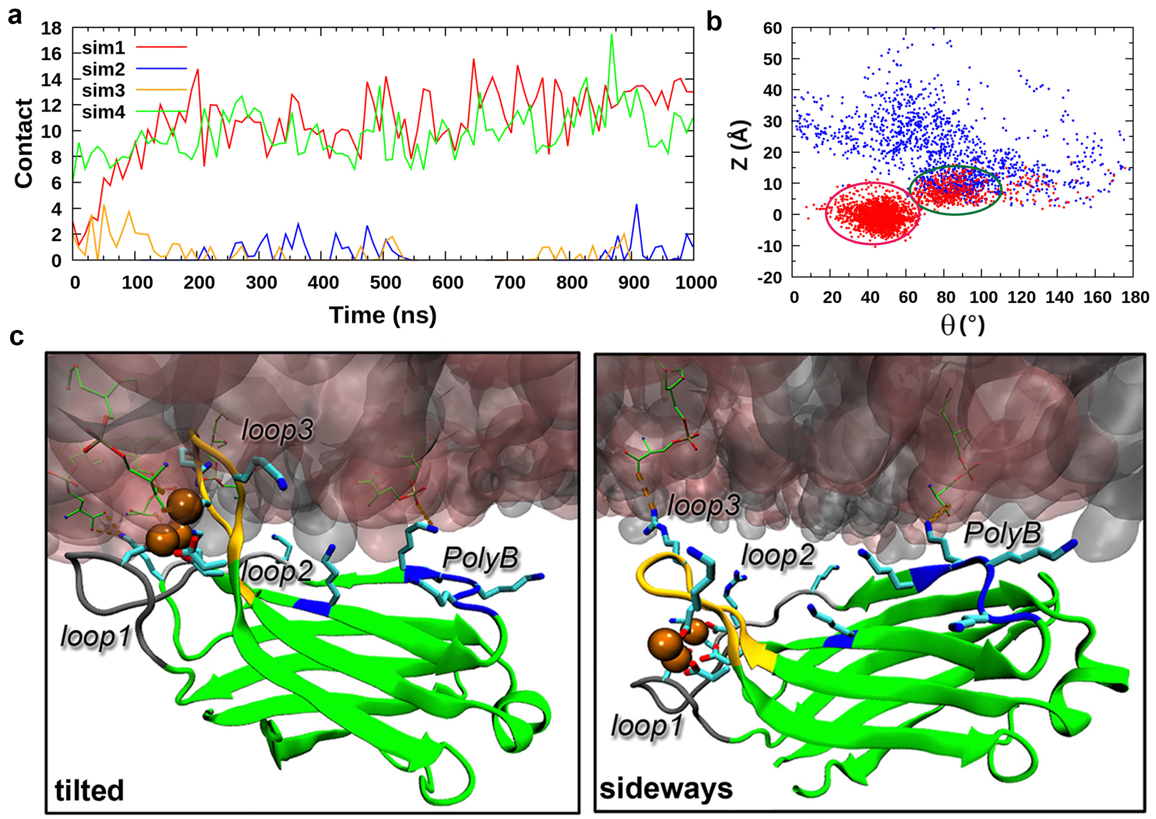


**Supplementary Figure 6. Membrane contacts and configurations of C2A in the FLΔC2B-VM simulations.** (a) Time traces of the total VM-contact numbers. (b) Scatter plot of Θ and Z. Red and blue dots are from snapshots where C2A contacts and does not contact VM, respectively; red and green ovals indicate ensembles in the tilted and sideways configurations, respectively. (c) C2B in the tilted (left panel) or sideways (right panel) configuration.
